# Comparison between an exact and a heuristic neural mass model with second order synapses

**DOI:** 10.1101/2022.06.15.496262

**Authors:** Pau Clusella, Elif Köksal-Ersöz, Jordi Garcia-Ojalvo, Giulio Ruffini

## Abstract

Neural mass models (NMMs) are designed to reproduce the collective dynamics of neuronal populations. A common framework for NMMs assumes heuristically that the output firing rate of a neural population can be described by a static nonlinear transfer function (NMM1). However, a recent exact mean-field theory for quadratic integrate-and-fire (QIF) neurons challenges this view by showing that the mean firing rate is not a static function of the neuronal state but follows two coupled non-linear differential equations (NMM2). Here we analyze and compare these two descriptions in the presence of second-order synaptic dynamics. First, we derive the mathematical equivalence between the two models in the infinitely slow synapse limit, i.e., we show that NMM1 is an approximation of NMM2 in this regime. Next, we evaluate the applicability of this limit in the context of realistic physiological parameter values by analyzing the dynamics of models with inhibitory or excitatory synapses. We show that NMM1 fails to reproduce important dynamical features of the exact model, such as the self-sustained oscillations of an inhibitory interneuron QIF network. Furthermore, in the exact model but not in the limit one, stimulation of a pyramidal cell population induces resonant oscillatory activity whose peak frequency and amplitude increase with the self-coupling gain and the external excitatory input. This may play a role in the enhanced response of densely connected networks to weak uniform inputs, such as the electric fields produced by non-invasive brain stimulation.

## 1 Introduction

Neural mass models (NMMs) provide a physiologically grounded description of the average synaptic activity and firing rate of neural populations (Wilson and Cowan, 1972; Lopes da Silva et al, 1974, 1976; Jansen et al, 1993; Jansen and Rit, 1995; Wendling et al, 2002). First developed in the 1970s, these models are increasingly used for both local and whole-brain modeling in, e.g., epilepsy (Wendling et al, 2002; Wendling and Chauvel, 2008; Jedynak et al, 2017) or Alzheimer’s disease (Pons et al, 2010; Stefanovski et al, 2019), and for understanding and optimizing the effects of transcranial electrical stimulation (tES) (Molaee-Ardekani et al, 2010; Merlet et al, 2013; Kunze et al, 2016; Ruffini et al, 2018; Sanchez-Todo et al, 2018). However, they are only partly derived from first principles. While the post-synaptic potential dynamics are inferred from data and can be grounded on diffusion physics (Destexhe et al, 1998; Pods et al, 2013; Ermentrout and Bard, 2010), the transfer function linking the mean population membrane potential with the corresponding firing rate (Freeman’s “wave-to-pulse” sigmoid function) rests on a weaker theoretical standing (Wilson and Cowan, 1972; Freeman, 1975; Kay, 2018; Eeckman and J, 1991). This results in a limited understanding on the range of applicability of the theory. For example, although models for the effects of an electric field at the single neuron are now available (Aberra et al, 2018; Galan, 2021), it is unclear how they should be used at the population-level representation.

In 2015, Montbrió, Pazó, and Roxin (MPR) (Montbrió et al, 2015) derived an exact mean-field theory for networks of quadratic integrate-and-fire (QIF) neurons, thereby connecting microscale neural mechanisms with mesoscopic brain activity. Within this framework, the response of a neural population is described by a low-dimensional system representing the dynamics of the firing rate and mean membrane potential. Therefore, the MPR equations can be seen to replace the usual static transfer sigmoid function with two differential equations grounded on the biophysics of the single neurons. Since then, the theory has been applied to cover increasingly complex formulations of the single-neuron activity, including time-delays (Pazó and Montbrió, 2016; Devalle et al, 2018; Ratas and Pyragas, 2018) dynamic synapses (Montbrió et al, 2015; Ratas and Pyragas, 2016; Devalle et al, 2017; Dumont and Gutkin, 2019; Coombes and Byrne, 2019; Byrne et al, 2020, 2022), gap-junctions (Laing, 2015; Pietras et al, 2019), stochastic fluctuations (Ratas and Pyragas, 2019; Goldobin et al, 2021; Clusella and Montbrió, 2022), asymmetric spikes (Montbrió and Pazó, 2020), sparse connectivity (di Volo and Torcini, 2018; Bi et al, 2021), and short-term plasticity (Taher et al, 2020, 2022).

In the limit of very slow synapses, the firing rate of the MPR formulation can be cast as a static function of the input currents, in the form of a population-wide *f-I* curve (Devalle et al, 2017). This function can be used to derive an NMM with exponentially decaying synapses, which fails to reproduce the dynamical behavior of the exact mean-field theory, highlighting the importance of the dynamical equations in the MPR model (Devalle et al, 2017). In fact, empirical evidence suggests that post-synaptic currents display rise and a decay time scales (Lopes da Silva et al, 1974; Jang et al, 2010). These type of synaptic dynamics can be modelled through a second-order linear equation, which forms the basis for many NMMs (see e.g. Lopes da Silva et al (1974); Jansen et al (1993); Wendling et al (2002)). This has been also noticed by other researchers, who have recently studied exact NMMs with second-order synapses (Coombes and Byrne, 2019; Byrne et al, 2020, 2022). However, a formal comparison between the MPR formalism with second-order synaptic dynamics and classical, heuristic NMMs has not yet been established.

In this paper we analyze the NMM that results from applying the mean-field theory to a population of QIF neurons with second-order equations for the synaptic dynamics. The resulting NMM, which we refer to as NMM2 in what follows, contains two relevant time scales: one for the post-synaptic activity and one for the membrane dynamics. These two time scales naturally bridge the Freeman “wave-to-pulse” function with the nonlinear dynamics of the firing rate. In particular, following Devalle et al (2017), we show that, in the limit of very slow synapses and external inputs, the mean membrane potential and firing rate dynamics become nearly stationary. This allows us to develop an analogous NMM with a static transfer function, which we will refer to as NMM1 for brevity. Next, we analyze the dynamics of the two models using physiological parameter values for the time constants, in order to assess the validity of the formal mapping. Bifurcation analysis of the two systems shows that the models are not equivalent, with NMM2 presenting a richer dynamical repertoire, including resonant responses to external stimulation in a population of pyramidal neurons, and self-sustained oscillatory states in inhibitory interneuron networks.

## 2 Models

### 2.1 NMM with static transfer function

Semi-empirical “lumped” NMMs where first developed in the early 1970s by Wilson and Cowan (Wilson and Cowan, 1972), Freeman (Freeman, 1972, 1975), and Lopes da Silva (Lopes da Silva et al, 1974). This framework is based on two key conceptual elements. The first one consists of the filtering effect of synaptic dynamics, which transforms the incoming activity (quantified by firing rate) into a mean membrane potential perturbation in the receiving population. The second element is a static transfer function that transduces the sum of the membrane perturbations from synapses and other sources into an output mean firing rate (see Grimbert and Faugeras (2006) for a nice introduction to the Jansen-Rit model). We next describe these two elements separately.

The synaptic filter is instantiated by a second-order linear equation coupling the mean firing rate of arriving signals *r* (in kHz) to the mean post-synaptic voltage perturbation *u* (in mV) (Grimbert and Faugeras, 2006; Ermentrout and Bard, 2010):

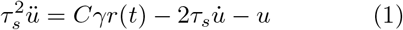

Here the parameter *τ_s_* sets the delay time scale (ms), *γ* characterizes the amplification factor in mV/kHz, and *C* is dimensionless and quantifies the average number of synapses per neuron in the receiving population. Upon inserting a single Dirac-delta-like pulse rate at time *t* = 0, the solution of (1) reads 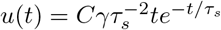 for a system initially at rest 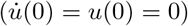. This model for PSPs activity is a commonly-used particular case of a more general formulation that considers different rise and decay times for the post-synaptic activity (Ermentrout and Bard, 2010).

The synaptic transmission equation needs to be complemented by a relationship between the level of excitation of a neural population and its firing rate, namely, a transfer function, Φ. Through the transfer function, each neuron population converts the sum of its input currents, *I*, to an output firing rate *r* in a non-linear manner, i.e., *r*(*t*) = Φ[*I*(*t*)]. Wilson and Cowan, and independently Freeman, proposed a sigmoid function as a simple model to capture the response of a neural mass to inputs, based on modeling insights and empirical observations (Wilson and Cowan, 1972; Freeman, 1975; Eeckman and J, 1991). A common form for the sigmoid function is

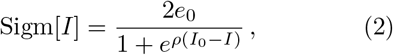

where *e*_0_ is the half-maximum firing rate of the neuronal population, *I*_0_ is the threshold value of the input (when the firing rate is *e*_0_), and *ρ* determines the slope of the sigmoid at that threshold. Beyond this sigmoid, transfer functions can be derived from specific neural models such as the leaky integrate-and-fire or the exponential integrate-and-fire, either analytically or numerically fitting simulation data, see e.g. Fourcaud-Trocmé et al (2003); Brunel and Hakim (2008); Pereira and Brunel (2018); Ostojic and Brunel (2011); Carlu et al (2020). In some studies, Φ is regarded as a function of mean membrane potential instead of the input current (Jansen and Rit, 1995; Wendling et al, 2002). Nonetheless, the relation between input current and mean voltage perturbation is often assumed to be linear, see for instance Ermentrout and Bard (2010). Therefore, the difference between both formulations might be relevant only in the case where the transfer function has been experimentally or numerically derived.

The form of the total input current in Eq. (2) will depend on the specific neuronal populations being considered, and on the interactions between them. In what follows we focus on a single population with recurrent feedback and external stimulation. Hence the total input current is given as the contribution of three independent sources,

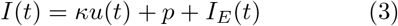

where *κ* is the recurrent conductance, *p* is a constant baseline input current, and *I_E_* stands for the effect of an electric field. Note that some previous studies do not use an explicit self-connectivity as an argument of the transfer function (see e.g. Grimbert and Faugeras (2006); Wendling et al (2002); Lopez-Sola et al (2021)). In the next section we show that the term *κu*(*t*) in (3) arises naturally in recurrent networks.

Finally, we rescale the postsynaptic voltage by defining *s* = *u*/(*Cγ*), and use the auxiliary variable *z* to write Eq. (1) as a system of two first-order differential equations. With those choices, the final closed formulation for the neural population dynamics reads

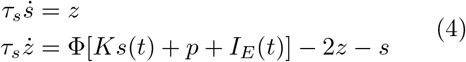

where *K* = *Cγκ*. We refer to this model in what follows as NMM1.

### 2.2 Quadratic integrate-and-fire neurons and NMM2

Consider a population of fully and uniformly connected QIF neurons indexed by *j* = 1,…, *N*. The membrane potential dynamics of a single neuron in the population, *U_j_*, is described by (Latham et al, 2000; Devalle et al, 2017)

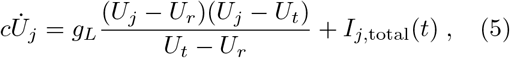

with *U_j_* being reset to *U*_reset_ when *U_j_* ≥ *U*_apex_. In this equation, *U_r_* and *U_t_* > *U_r_* represent the resting and threshold potentials of the neuron (mV), *I*_*j*,total_ the input current (*μA*), *c* the membrane capacitance (*μF*), and *g_L_* is the leak conductance (mS). If unperturbed, the neuron membrane potential tends to the resting state value *U_r_*. In the presence of input current, the membrane potential of the neuron *U_j_* can grow and surpass the threshold potential *U_t_*, at which point the neuron emits a spike. An action potential is produced when *U_j_* reaches a certain apex value *U*_apex_ > *U_t_*, at which point *U_j_* is reset to *U*_reset_.

The total input current of neuron *j* is

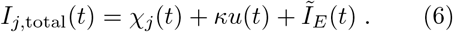

The first term in this expression, *χ_j_*(*t*), corresponds to a Cauchy white noise with median 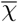 and half-width at half-maximum Γ (see Clusella and Montbrió (2022)). The second term, *κu*(*t*), represents the mean synaptic transmission from other neurons *u*(*t*), with coupling strength κ. As in NMM1, we assume that *u*(*t*) follows Eq. (1). However, in this case, the firing rate is determined self-consistently from the population dynamics as

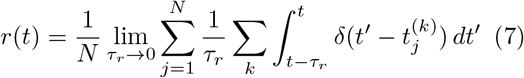

where 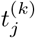 is the time of the *k*th spike of neuron *j*, and the spike duration time *τ_r_* needs to assume small finite values in numerical simulations. Finally, *Ĩ_E_*(*t*) can represent both a common external current from other neural populations, or the effect of an electric field. In the case of an electric field, the current can be approximated by 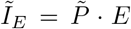, where 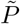 is the dipole conductance term in the spherical harmonic expansion of the response of the neuron to an external, uniform electric field (Galan, 2021). This is a good approximation if the neuron is in a subthreshold, linear regime and the field is weak, and can be computed using realistic neuron compartment models. We assume here for simplicity that all the QIF neurons in the population are equally oriented with respect to the electric field (this could be generalized to a statistical dipole distribution).

In order to analyze the dynamics of the model it is convenient to cast it in a reduced form. Following Devalle et al (2017), we define the new variables

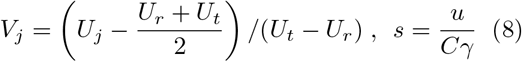

and redefine the system parameters (all dimensionless except for *τ_m_*) as

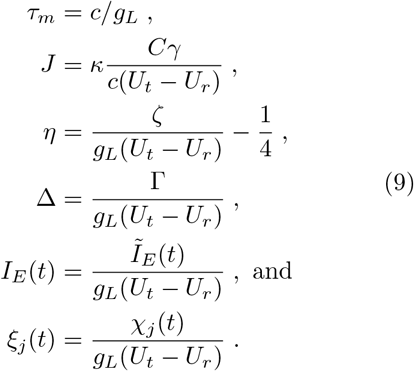

With these transformations, the QIF model can be written as

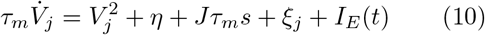

with the synaptic dynamics given by

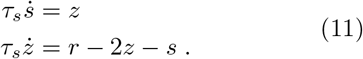

These transformations express the QIF variables and parameters with respect to reference values of time (*c/g_L_*), voltage (*U_t_* – *U_r_*), and current (*g_L_*(*U_t_* – *U_r_*)). In the new formulation, the only dimensional quantities have units of time (*τ_m_* and *τ_s_*, in ms) or frequency (*r* and *s*, in kHz). It is important to keep in mind these changes when dealing with multiple interacting populations involving different parameters, and also when using empirical measurements to determine specific parameter values.

#### 2.2.1 Exact mean-field equations with second order synapses (NMM2)

Starting from Eq. (10), Montbrió et al (2015) derived an effective theory of fully connected QIF neurons in the large *N* limit. Initially, the theory was restricted to deterministic neurons with Lorentzian distributed currents. Recently it has also been shown to apply to neurons under the influence of Cauchy white noise, a type of Lévy process that renders the problem analytically tractable Clusella and Montbrió (2022). In any case, the macroscopic activity of a population of neurons given by Eq. (10) can be characterized by the probability of finding a neuron with membrane potential *V* at time *t*, *P*(*V*, *t*). In the limit of infinite number of neurons (*N* → ∞), the time evolution of such probability density is given by a Generalized Master Equation (GME). Assuming that the reset and threshold potentials for single neurons are set to *V*_apex_ = – *V*_reset_ = ∞, the GME can be solved by considering that *P* has a Lorentzian shape in terms of a time-depending mean membrane potential *v*(*t*) and mean firing rate *r*(*t*),

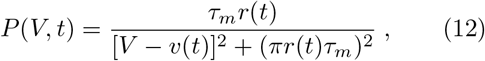

with

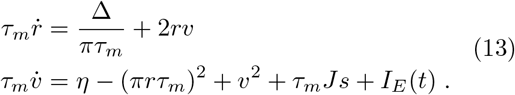

Together with the synaptic dynamics (11), these equations describe an exact NMM, which we refer to as NMM2.

## 3 Slow and fast synapse dynamics limits

### 3.1 Slow synapse limit and map to NMM1

Comparing the formulations of the semi-empirical model NMM1 (4) and the exact mean-field model NMM2 (13) one readily observes that the latter can be interpreted as an extension of the former. The synaptic dynamics are given by the same equations in both models, yet in NMM2 the firing rate *r* is not a static function of the input currents, but a system variable. Moreover, NMM2 includes the dynamical effect of the mean membrane potential, *v*, which in the classical framework is assumed to be directly related with the post-synaptic potential.

Devalle et al (2017) showed that in a model with exponentially decaying (i.e. first-order) synapses, the firing rate can be expressed as a transfer function in the limit of slowly decaying synapses. Their work follows from previous results showing that, in class 1 neurons, the slow synaptic limit allows one to derive firing rate equations for the population dynamics (Ermentrout, 1994). Here we revisit the same steps to show that NMM2 can be formally mapped to a NMM1 form. We perform such derivation in the absence of external inputs (*I_E_*(*t*) = 0).

Let us rescale time in Eq. (13) to units of *τ_s_*, and the rate variables to units of 1/*τ_m_* using

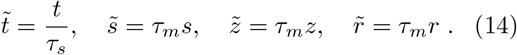

Additionally we define *ϵ* = *τ_m_/τ_s_*. Then the NMM2 model (Eqs. (11) and (13)) reads

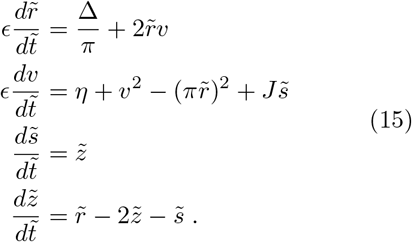

Taking now *ϵ* → 0 (*τ_s_* → ∞), the equations for 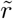 and *v* become quasi-stationary in the slow time scale, i.e.

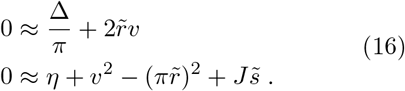

The solution of these equations is given by 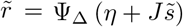, where

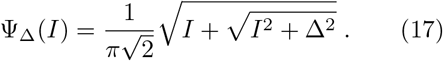

This is the transfer function of the QIF model, which relates input currents to the output firing rate. Thus, in the limit *ϵ* → 0 system (15) formally reduces to the NMM1 formulation, Eq. (4).

In following sections we study to what extent this equivalence remains valid for finite ratios of *τ_m_/τ_s_*. To that end, it is convenient to recast the analogy between NMM1 and NMM2 in terms of the non-rescaled quantities, which corresponds to using

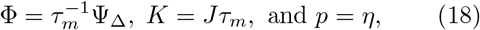

in Eq. (4).

In Fig 1 we fit the parameters of the sigmoid function to Ψ_Δ_ for Δ = 1. Despite the sudden sharp increase of both functions, there is an important qualitative difference: the *f-I* curve of the QIF model does not saturate for *I* → ∞. Other transfer functions derived from neural models share a similar non-bounded behavior (Fourcaud-Trocmé et al, 2003; Carlu et al, 2020). This reflects the continued increase of firing activity with increase input, which has been reported in experimental studies (Rauch et al, 2003).

**Figure 1.**
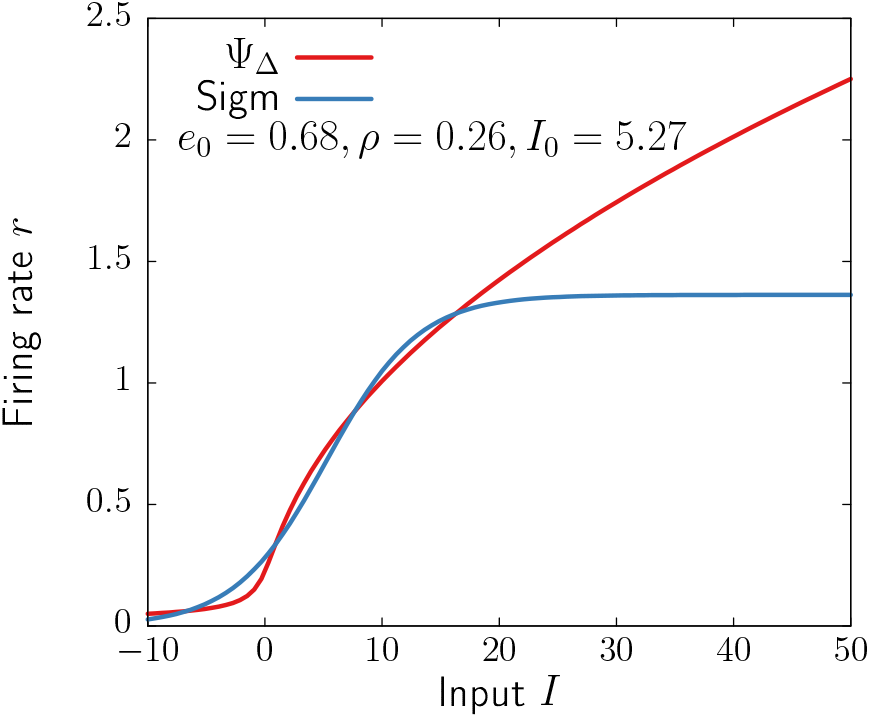
Transfer function of the QIF network (17) with Δ = 1 and the sigmoid (2) with parameters fitted to Φ_Δ_.

### 3.2 Fast synapse limit

To explore the fast synapse limit it is convenient to rescale time as 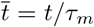 in the NMM2 equations. In this new frame, and defining *δ*:= *τ_s_/τ_m_* = 1/*ϵ*, the system reads

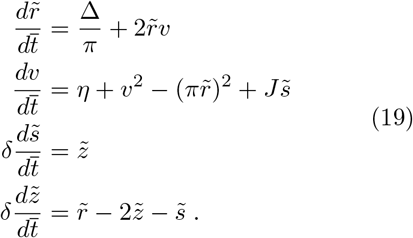

where 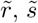 and 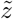 are the rescaled variables defined in (14). With the algebraic conditions in the fast synapse limit, *δ* → 0 (*τ_s_* → 0), Eq. (19) is reduced to

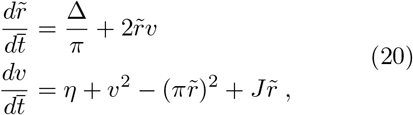

where we have used that 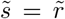 as given by the synaptic equations. This is the model with instantaneous synapses analyzed by Montbrió et al (2015), who showed that the *η–J* phase diagram has three qualitatively distinct regions in the presence of a constant input: a single stable node corresponding to a low-activity state, a single stable focus (spiral) generally corresponding to a high-activity state, and a region of bistability between a low activity steady state and a regime of asynchronous persistent firing.

## 4 System dynamics

In the previous section we have shown that NMM2 can be mapped to NMM1 in the limit of slow synapses (*τ_m_/τ_s_* → 0), using the scaling relations (18). However, physiological values for the time constants might not be consistent with this limit. Table 1 shows reference values for *τ_m_* and *τ_s_* corresponding to different neuron types and their corresponding neurotransmitters obtained from experimental studies. Notice that, in practice, such values also depend on the electrical and morphological properties of the neurons, and pre- and post-neuron types. Such level of detail requires the use of conductance-based compartmental models, a further step in mathematical complexity that is out of the scope of this paper. Therefore, we take the values in Table 1 as coarse-grained quantities that properly reflect the time scales in point neuron models such as the QIF (10) (for a more detailed discussion see Section 5). In order to study to what extend these non-vanishing values of *τ_m_/τ_s_* break down the equivalence between the two models, in this section we analyze and compare the dynamics of a single neural population with recurrent connectivity described by both NMM1 (Eq. (4) with Eq. (18)) and NMM2 (Eqs. (11) and (13)).

**Table 1.** Values for the membrane time constants *τ_m_* and postsynaptic currents *τ_s_*, for Pyramidal neurons (Pyr), parvalbumin-positive (PV+), and neurogliaform cells (NGFC) (Neske et al, 2015; Zaitsev et al, 2012; Povysheva et al, 2007; Avermann et al, 2012; Oláh et al, 2007; Seay et al, 2020; Karnani et al, 2016; Bacci et al, 2003; Deleuze et al, 2019). Notice that, in general, the synaptic time-constant should depend on the neurotransmitter, and the pre- and post-synaptic cells. Since we only consider self-coupled populations, we do not specify time-constants for transmission across populations of different types.

| Neuron type | Neurotransmitter | $\tau_m$ (ms) | $\tau_s$ (ms) | $\tau_m/\tau_s$ |
| --- | --- | --- | --- | --- |
| Pyr | Glutamate | 15 | 10 | 1.5 |
| PV+ | GABA | 7.5 | 2 | 3.75 |
| NGFC | GABA | 11 | 20 | 0.55 |

The first step is to identify the steady states of the system. Since we derived the transfer function (17) by assuming the *r* and *v* variables of NMM2 to be nearly stationary, the fixed points of both models coincide and are given by

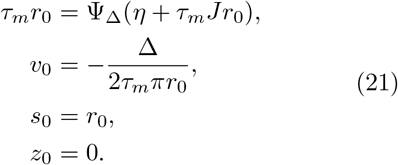

Moreover, *r*_0_ and *v*_0_ are the equilibrium points of the two-dimensional system analyzed by Montbrió et al (2015). Notice that the only relevant parameters for the determination of the fixed points are *J*, *η*, and Δ. The time constant *τ_m_* only acts as a multiplicative factor of *r*_0_ (and *s*_0_), and *τ_s_* does not enter into the expressions of the steady states.

Even though the steady states of the three models (NMM1, NMM2, and the original system of Montbrió et al (2015)) are the same, their stability properties might be different, as we now attempt to elucidate. The eigenvalues controlling the stability of the fixed points in NMM1 are

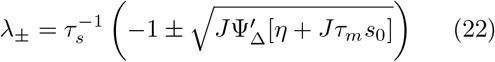

A similar closed expression for NMM2 is complicated to obtain and, in any case, there are no explicit expressions for the steady states. Thus, we use in what follows the numerical continuation software AUTO-07p (Doedel et al, 2007) to obtain the corresponding bifurcation diagrams. We analyze separately the dynamics of excitatory (*J* > 0) and inhibitory (*J* < 0) neuron populations in the two NMM models.

### 4.1 Pyramidal neurons

We start by analysing the dynamics of NMM1 in the case of excitatory coupling (*J* > 0), by fixing Δ = 1 and varying *η* and *J*. Following Table 1, we set *τ_m_* = 15 ms and *τ_s_* = 10 ms. Since the NMM1 eigenvalues (22) are real for *J* > 0, the fixed points do not display resonant behavior, i.e., they are either stable or unstable nodes. For positive baseline input *η*, only a single fixed point exists irrespective of the value of the coupling *J*. In contrast, a large region of bistability bounded by two saddle-node (SN) bifurcations emerges for negative *η*. The green curves in Fig. 2(a) show the two SN bifurcations, which merge in a cusp close to the origin of parameter space. Within the region bounded by the curves (green shaded region), a low-activity and a high-activity state coexist, separated by a third unstable fixed point. Figure 2(b) displays, for instance, the stationary firing rate as a function of *η* for *J* = 40. The values of the time constants *τ_m_* and *τ_s_* do not affect the bistability region. However, the noise amplitude Δ does have an effect: as shown in Appendix A, NMM1 admits a parameter reduction that expresses all parameters and variables as functions of Δ. Accordingly, the effects of modifying Δ on the stability of the fixed points are analogous to rescaling *η* → *η*/Δ and 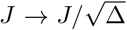 (see also Montbrió et al (2015)). Therefore, the bistable region shrinks in the (*η*, *J*) parameter space as the noise amplitude increases.

**Figure 2.**
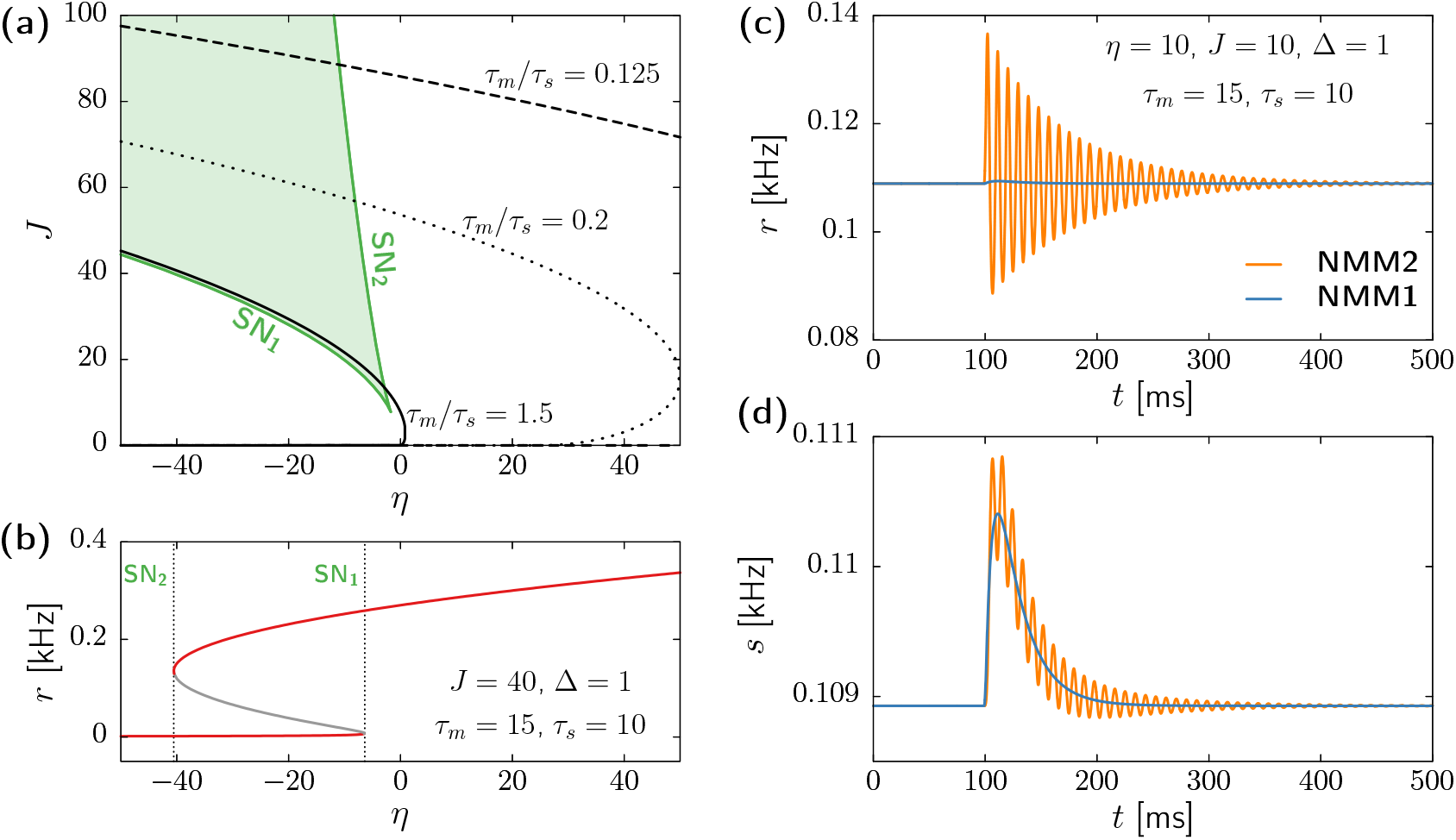
Dynamics of a population of pyramidal neurons described by NMM1 and NMM2. (a) Saddle-node bifurcations SN_1_ and SN_2_ (green curves) limiting the region of bistability (light-green region), and node-focus boundary for three different values of *τ_m_/τ_s_* in NMM2 (solid, dotted and dashed black lines). (b) Steady-state value of the firing rate as a function *η*, for fixed *J* = 40, Δ = 1, *τ_m_* = 15 ms, and *τ_s_* = 10 ms. The stable steady state branches are colored in red, and the unstable steady state branch in grey. Dashed vertical lines indicate the SN1 and SN2 bifurcation points (cf. panel a). (c,d) Time evolution of the firing rate *r* (c) and synaptic variable *s* (d) for NMM1 (blue) and NMM2 (orange), for *η* = 10, *J* = 10, Δ = 1, *τ_m_* = 15ms, and *τ_s_* = 10 ms. Initial conditions are at steady state, and a 1 ms-long pulse of *I_E_*(*t*) = 10 is applied at t = 100 ms.

Since the fixed points of NMM1 and NMM2 coincide, these two branches of SN bifurcations also exist in NMM2. Moreover, no other bifurcations arise, thus the diagrams depicted in Figs. 2(a,b) also hold for the exact model. However, there is an important difference regarding the relaxation dynamics towards the fixed points: While in NMM1 the steady states are always nodes, in NMM2 trajectories near the high-activity state might display transient oscillatory behavior. Figures 2(c,d) display, for instance, time series obtained from simulations of NMM1 (blue) and NMM2 (orange) starting at the fixed point, and receiving a small pulse applied at *t* = 100 ms. Not only NMM2 displays an oscillatory response, but also the effect of the perturbation in the firing rate is much larger in NMM2 than in NMM1.

Such resonant behavior of NMM2 corresponds to the two dominant eigenvalues of the high-activity fixed point (those with largest real part) being complex conjugates of each other. The black curves in Fig 2(a) show the boundary line at which those two eigenvalues change from real (below the curves) to complex (above the curves). For physiological values of *τ_m_* and *τ_s_* (continuous black line) this node-focus line remains very similar to that of the model with instantaneous synapses studied in Montbrió et al (2015). Reducing the ratio *τ_m_/τ_s_* changes this situation. As shown by the dotted and dashed curves in Fig. 2(a), as we approach the slow synaptic limit (*τ_m_/τ_s_* → 0) the resonant region (where the dominant eigenvalues are complex) requires increasingly larger values of *η* and Δ, vanishing for small enough ratio *τ_m_/τ_s_*. Hence, as expected from the time scale analysis of Section 3, the dynamics of NMM2 can be faithfully reproduced by NMM1 in this limit. However, the equivalence cannot be extrapolated to physiological parameter values.

### 4.2 Interneurons

Here we consider a population of GABAergic interneurons with self-recurrent inhibitory coupling (*J* < 0). In particular, we focus on parvalbumin-postive (PV+) fast spiking neurons, which play a major role in the generation of fast collective brain oscillations (Bartos et al, 2002, 2007; Cardin et al, 2009; Tiesinga and Sejnowski, 2009). We thus set *τ_m_* = 7.5 and *τ_s_* = 2 ms, following Table 1.

In this case, the NMM1 dynamics are rather simple: there is a single fixed point that remains stable, with a pair of complex conjugate eigenvalues (see Eq (22)). Therefore, the transient dynamics do display resonant behavior upon external perturbation. Nonetheless, no self-sustained oscillations emerge.

In the NMM2, however, the unique fixed point might lose stability for *η* > 0 through a supercritical Hopf bifurcation (HB+, see blue curve in Fig. 3(a)). This transition gives rise to a large region of fast oscillatory activity, corresponding to the so-called interneuron-gamma (ING) oscillations (Whittington et al, 1995; Traub et al, 1998; Whittington et al, 2000; Bartos et al, 2007; Buzsáki and Wang, 2012). An example of this regime is shown in Figs. 3(b,c), using both NMM2 as well as microscopic simulations of a QIF network as defined by Eq. (10).

**Figure 3.**
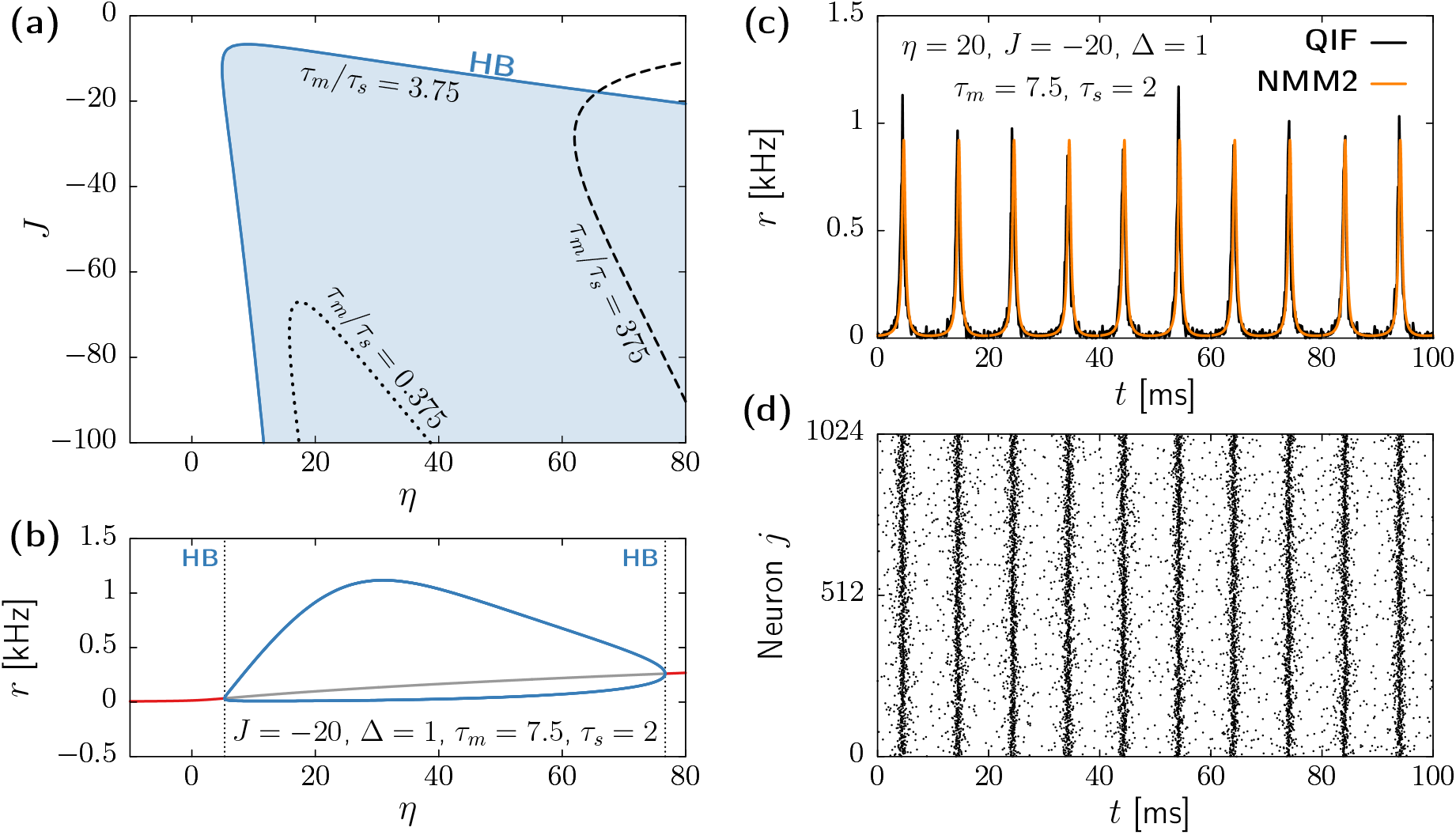
Dynamics of a population of parvalbumin-positive interneurons described by NMM1 and NMM2. (a) Supercritical Hopf bifurcation signaling the onset of oscillatory activity for *τ_m_* = 7.5 ms, *τ_s_* =2 *ms* (blue curve), *τ_m_* = 7.5 ms, *τ_s_* = 20 ms (dotted black curve), and *τ_m_* = 7.5 ms, *τ_s_* = 0.02 ms (dashed black curve). The blue-shaded region indicates stable limit-cycle behavior for *τ_m_* = 7.5 ms, *τ_s_* = 2 ms. (b) Steady-state values of the firing rate as a function of the input *η*, for fixed *J* = −20, Δ = 1, *τ_m_* = 7.5 ms, and *τ_s_* = 2 ms. The red line represents the stable steady state, the grey line the unstable steady state, and the blue lines the maxima and minima of the stable limit-cycle. Dashed vertical lines indicate the location of supercritical Hopf bifurcations (cf panel a). (c) Time evolution of the firing rate *r* for an inhibitory population at the oscillatory state (*η* = 20, *J* = −20, Δ = 1, *τ_m_* = 7.5 ms, and *τ_s_* = 2 ms) obtained from integrating the a network with *N* = 1024 QIF neurons (10) (black) and from the NMM2 (13) (orange). (d) Raster plot of the spiking times in the simulation of the QIF network corresponding to panel (c). Simulations of QIF network were performed with *V*_apex_ = – *V*_reset_ = 100 using Euler-Maruyama integration with *dt* = 10^-3^ ms. The firing rate *r* was computed using Eq. (7) with *τ_r_* = 10^-2^ ms.

According to the ING mechanism, oscillations emerge due to a phase lag between two opposite influences: the noisy excitatory driving (controlled by *η* and Δ) and the strong inhibitory feedback from the recurrent connections (controlled by *J*). In NMM2, the dephasing between these two forces stems from the implicit delay caused by the synaptic dynamics. Hence, the ratio between membrane and synaptic characteristic times, *τ_m_* and *τ_s_*, has a fundamental role in the generation of ING oscillations. The blue region depicted in Fig. 3(a) corresponds to time scales of PV+ neurons, *τ_m_* = 7.5 and *τ_s_* = 2. In this case the oscillation frequency is in the gamma range (40-200Hz). However, by decreasing the parameter ratio *τ_m_/τ_s_* the Hopf bifurcation becomes elusive, as the oscillatory region shrinks, and oscillations require stronger inhibitory feedback (see black dotted curve in Fig. 3(a)). Similarly, by increasing *τ_m_/τ_s_* the ING activity also fades, as larger inputs **η** are required to produce oscillatory activity (see black dashed curve in Fig 3(a)). As showed in the previous section, the two limits of *τ_m_/τ_s_* coincide with NMM1 and the model analyzed in Montbrió et al (2015). The results presented above show that the membrane and synaptic dynamics are required to have comparable time scales, in order to generate oscillatory activity in NMM2.

### 4.3 Network-enhanced resonance in excitatory populations

The bifurcation analysis of Section 4.1 reveals that a single population of excitatory neurons does not display self-sustained oscillations in neither NMM1 nor NMM2. This is expected, as excitation alone is known to be usually insufficient for the emergence of collective rhythms (Van Vreeswijk et al, 1994). However, in NMM2, the high-active steady state corresponds to a stable focus in a large region of the parameter space. In this section we exploit this resonant behavior, inspired by the oscillatory response of a population of pyramidal neurons subject to tACS stimulation. We thus consider the NMM2 model with *τ_m_* = 15 ms and *τ_s_* = 10 ms injected with a current

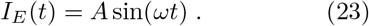

We expect to induce oscillatory activity if *ω* is close to the resonant frequency of the system, given by *ν*:= Im[λ], where λ is the fixed point eigenvalue with largest real part.

Figures 4(a,b) display heatmaps of the standard deviation of the firing rate, *σ*, obtained by stimulating the stable focus of NMM2 at different frequencies *ω* and amplitudes *A*. For weak baseline input *η* (Fig. 4(a)), the amplitude of the system displays a large tongue-shaped region, with a few additional narrow tongues at smaller frequencies. The main tongue is centered at the resonant frequency *ω* ≃ *ν* (see grey vertical dashed line) and corresponds to entrainment at the driving rhythm, whereas secondary tongues correspond to entrainment at higher harmonics. Increasing the external input *η* (Fig. 4(b)) causes the system to resonate at larger frequencies, and shrinks the region of amplification of the applied stimulus. Despite the similitude with the usual Arnold tongues that characterize driven oscillatory systems, we recall that here we are inducing oscillatory activity in an otherwise stationary system. Hence, even if small in amplitude, there is always an oscillatory response at some harmonic of the driving frequency.

**Figure 4.**
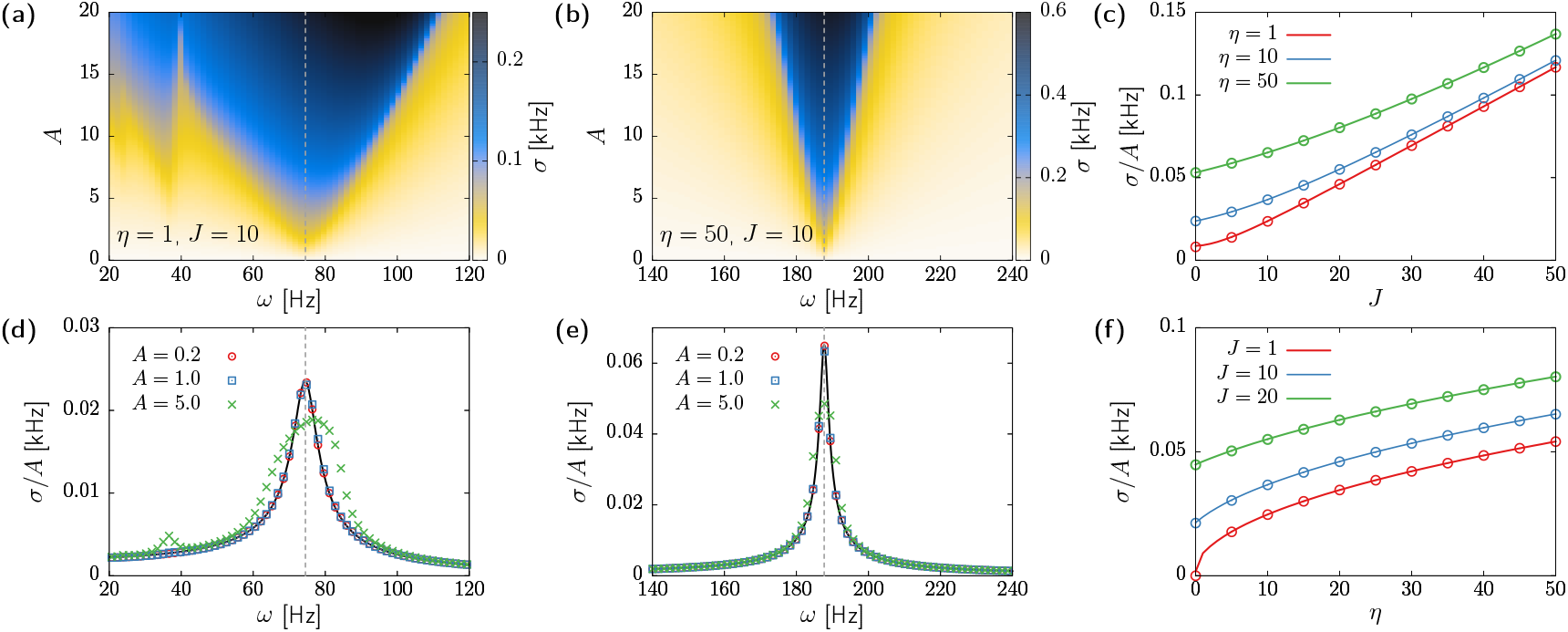
Effects of tACs stimulation, Eq. (23), in a population pyramidal neurons given by NMM2 (13). (a,b) Heatmaps of the standard deviation of *r* displaying Arnold tongues for *η* =1 (panel (a)) and *η* = 50 (panel(b)). The rest of system parameters are *J* = 10, Δ = 1, *τ_m_* = 15 ms, and *τ_s_* = 10 ms. (c) Normalized amplitude *σ/A* obtained by stimulating the population at its resonant frequency *ν*, for increasing values of the coupling strength. Continuous lines correspond to analytical results (Eq. (24)), and circles correspond to numerical simulations. (d,e) Normalized amplification *σ/A* corresponding to the same parameters of panels (a) and (b), respectively. Symbols correspond to the numerical simulations reported in (a) and (b). The black continuous lines correspond to Eq. (24). (f) Normalized amplitude *σ/A* at the resonant frequency *ν* upon increasing the external input *η*. Lines correspond to Eq. (24) and symbols to numerical simulations. In all panels, periodic stimulation has been simulated for 2 seconds after letting the system relax to the fixed point for 1 second. The reported values for the standard deviation *σ* correspond only to the last 1 s of stimulation, in order to avoid capturing transient effects.

Electric stimulation protocols usually achieve large effects even when the amplitudes of the oscillatory input signal are small. We thus investigate the effect of weak stimuli through a perturbative analysis for 0 < *A* ≪ 1. Upon expanding the NMM2 equations close to the fixed point and solving the resulting linear system, we obtain the amplitude response as a function of the driving frequency:

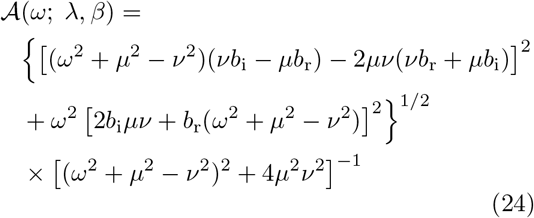

where λ = *μ* + *iν* is the fixed point eigenvalue with largest real part, and *b* = *b_r_* + *ib_i_* is the associated amplitude component (see Appendix B for the mathematical details). These two complex quantities can be obtained by numerically computing the eigenvalues and eigenvectors of the system Jacobian. The black curves in Fig 4(d) and (e) illustrate the validity of the analytical expression when compared with numerical results (colored symbols). Overall, the perturbative analysis provides a good approximation for *A* < 1, showing that, at this stage, *ω* = *ν* provides the maximal amplification.

Finally, we use these results to investigate the effect of the system parameters *J* and *η* to the amplitude response of the neural mass. Figs. 4(b,c) show numerical (open circles) and analytical (lines) results obtained using the optimal stimulation protocol *ω* = *ν* with *A* = 0.1 for different values of *J* and *η*. Overall, the oscillation amplitude of the system shows a supra-linear increase with *J*, and a sublinear increase with *η*. These results illustrate the importance of self-connectivity in tACs stimulation, and can potentially explain the effectiveness of these protocols in spite of the weakness of the applied electric field. Since we only considered driving of an excitatory population, the associated resonant frequencies can be quite large (up to 400Hz for *η* = 50 and *J* = 50), which calls for future investigations to analyze the combined effect of tACs in networks with excitation-inhibition balance.

## 5 Conclusions

For decades, NMMs have been built up on the basis of a simple framework that combines the linear dynamics of synaptic activation with a nonlinear static transfer function linking neural activity (firing rate) to excitability (Wilson and Cowan, 1972; Freeman, 1975; Lopes da Silva et al, 1974). This view has been sustained by empirical observations and heuristic assumptions underlying neural activity. Models based on this framework have been used to explain the mechanisms behind neural oscillations (Lopes da Silva et al, 1974; Freeman, 1987; Jansen and Rit, 1995; Wendling et al, 2002), and, more recently, to create large-scale brain models to address the treatment of neurophatologies by means of electrical stimulation (Kunze et al, 2016; Sanchez-Todo et al, 2018; Forrester et al, 2020).

Further theoretical efforts have provided more sophisticated tools to model the dynamics of neural populations, by deriving transfer functions from specific single-cell models (Gerstner, 1995; Brunel and Hakim, 2008; Ostojic and Brunel, 2011; Carlu et al, 2020), add adaptation mechanisms (Augustin et al, 2017), or finite size-effects (Benayoun et al, 2010; Buice et al, 2010). In this context, exact NMMs (also known as Next-Generation NMMs) pave a new road to directly relate single neuron dynamics with mesoscopic activity (Montbrió et al, 2015). Understanding how this novel framework relates to previous semi-empirical models should allow us to validate the range of applicability of classical NMMs.

Here we have studied a neural mass with second-order synapses, similar to the one studied in recent works (Coombes and Byrne, 2019; Byrne et al, 2020, 2022). The model naturally links the dynamical firing rate dynamics derived by Montbrió et al (2015) with the typical linear filtering representing synaptic transmission that is used in heuristic NMMs. Following Ermentrout (1994) and Devalle et al (2017) we show that, in the slow-synapse limit and in the absence of time-varying inputs, the exact model can be formally mapped to a simpler formalism with a static transfer function. However, we find that the range of validity of this relationship is beyond the physiological values of the model parameters. An analysis of the dynamics using realistic values of the time constants illustrates the fact that fundamental properties, such as the resonant behavior of excitatory populations and the interneuron-gamma oscillatory dynamics of PV+ neurons, cannot be captured by a traditional formulation of the model.

Despite the exact mean-field theory leading to NMM2 is a major step forward on the development of realistic mesoscale models for neural activity, the QIF neuron is a simplified model with some limitations. For instance, here we have employed non-refractory neurons, for which increasing input currents always lead to an increase of the firing rate. Future studies should address the role of a refractive period on the emerging rhythms and stimulation effects of exact NMMs. This could lead to a more realistic saturating shape of the QIF transfer function (Fig. 1). Additionally, further considerations may need to be taken into account in order to translate experimental observations to the model. In particular, the synapse time constants reported in Table 1 should reflect the delay and filtering associated with current transmission from input site to soma. This is not trivial to measure experimentally, and it can change considerably depending on synapse location, morphology, the number of simultaneously activated spine synapses (Eyal et al, 2018), and electrical properties (Koch and Segev, 2003), which are not accounted by the QIF neuron, but can be estimated using realistic compartment models (Agmon-Snir and Segev, 1993). Besides, the QIF model is an approximation of type-I excitable neurons, with type-II having a completely different firing pattern and *f-I* curve.

An important application of the exact meanfield theory is in the context of transcranial electrical stimulation. Several decades of research suggest that weak electric fields influence neural processing (Ruffini et al, 2020). In tES, the electric field generated on the cortex is of the order of 1 V/m, which is known to produce a sub-mV membrane perturbation (Bikson et al, 2004; Ruffini et al, 2013; Aberra et al, 2018). Yet, the applied field is mesoscopic in nature and is applied during long periods, with a spatial scale of several centimeters and temporal scales of thousands of seconds. Hence, a long-standing question in the field is how networks of neurons process spatially uniform weak inputs that barely affect a single neuron, but produce measurable effects in population recordings. By means of the exact mean-field model, we have shown that the sensitivity of the single population to such a weak alternating electric field can be modulated by the intrinsic self-connectivity and the external tonic input of the neural population in a population of excitatory neurons. Importantly, such resonant behavior cannot be captured by heuristic NMMs with static transfer functions.

For the physiologically-inspired parameter values chosen in this study, the amplification effects on excitatory neurons appear to be weaker than those observed experimentally. We may conjecture that certain neuronal populations may be in states near criticality, i.e. close to the bifurcation points in the NMM2 model (Chialvo, 2004; Carhart-Harris, 2018; Vázquez-Rodríguez et al, 2017; Zimmern, 2020; Ruffini and Lopez-Sola, 2022). This would apply, for example, to inhibitory populations, which display a Hopf bifurcation where a state near the critical point will display arbitrarily large amplified sensitivity to weak but uniform perturbations applied over long time scales. Since electric fields are expected to couple more strongly to excitatory cells, this case should be studied in the context of a multi-population NMM2, with excitatory cells relaying the electric field perturbation. Exact NMMs provide an appropriate tool to investigate this behavior, as well as the effects of non-homogeneous electrical fields—which we leave to future studies.

## Acknowledgments

This work has received funding from the European Research Council (ERC Synergy) under the European Union’s Horizon 2020 research and innovation programme (grant agreement No 855109) and from the Future and Emerging Technologies Programme (FET) under the European Union’s Horizon 2020 research and innovation programme (grant agreement No 101017716). J.G.-O. is supported by the Spanish Ministry of Science and Innovation and FEDER (grant PGC2018-101251-B-I00), by the “Maria de Maeztu” Programme for Units of Excellence in R&D (grant CEX2018-000792-M), and by the Generalitat de Catalunya (ICREA Academia programme).

## Appendix A Parameter reduction in NMM1 and NMM2

A common way to simplify the analysis of dynamical systems such as NMM1 (Eq. 4) and NMM2 (Eqs. 11,13) is through parameter reduction. While this can be achieved in different ways, here we choose, following Montbrió et al (2015), to rescale the system parameters as follows:

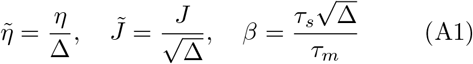

We then define the new variables

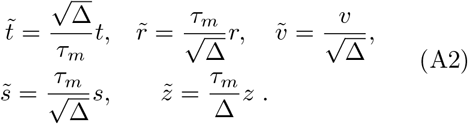

With these definitions, and together with the equivalence relation (18), NMM1 takes the form:

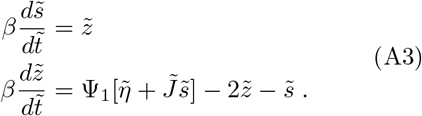

Similarly, the NMM2 model (Eqs. 11,13) becomes

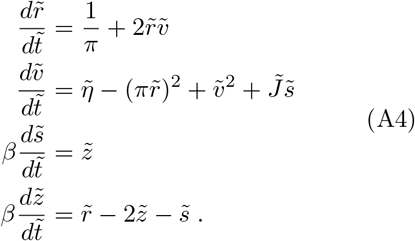

These reduced systems reveal that the dynamics of both models are controlled only by the three effective parameters 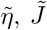, and *β*. In particular, the effects of changing Δ in the attractors of the system can be achieved by appropriately modifying the other parameters. Also, this reduction makes explicit that the bifurcations of the system do not depend on specific values of *τ_m_* and *τ_s_*, but only on their ratio. Notice, however, that in (A4) all parameters and variables, including time, become adimensional. This contrasts with the formulation used throughout the paper (Eqs. (13)), where time has units of miliseconds.

## Appendix B Analysis of a weakly periodically forced system

Here we present the results on weakly periodically perturbed systems used to investigate the response of NMM2 to periodic stimulation in section 4.3. Although we have a specific system in mind, we consider a general setup for simplicity. Let 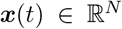 be an *N*-dimensional state vector, with time evolution given by the autonomous nonlinear system

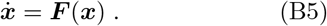

In NMM2, ***F*** follows Eqs. (11,13), and the state vector reads ***x*** = (*r*, *v*, *s*, *z*)^*T*^. Let ***x***^(0)^ be a stable fixed point of the system and ***J*** = ***J***(***x***^(0)^) the corresponding Jacobian. We consider a periodic forcing acting on Eq. (B5),

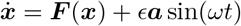

where *ϵ* is a weak coupling 0 < *ϵ* ≪ 1 and 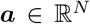 is a normalized vector for the distribution of the forcing across the system variables. For instance, in the case considered in the paper ***a*** = (0, 1, 0, 0)^*T*^, since the periodic driving acts only on the mean membrane potential.

Let ***x*** = ***x***^(0)^ + *ϵ**δx***. Since *ϵ* ≪ 1 we can linearize close to the fixed point, ***x***^(0)^, to obtain

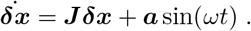

Let ***S*** be the matrix of eigenvectors of ***J***, and **Λ** the diagonal matrix of eigenvalues, so that ***S***^-1^***JS*** = **Λ**. The coordinates of the perturbation vector in the basis defined by the Jacobian eigenvalues read ***α***:= ***S***^-1^***δx***. Therefore,

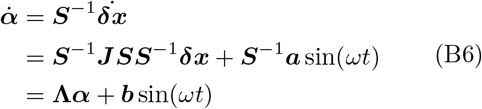

where ***b***:= ***S***^-1^***a***, i.e., the coordinates of ***a*** in the basis defined by ***S***.

Since **Λ** is a diagonal matrix, Eq. (B6) can be written in scalar form for each *α_j_* in complex space as

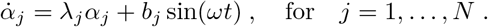

In what follows we drop the subindices *j* for simplicity. The solution of each of the linear systems for *α* read

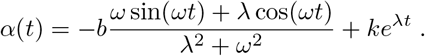

with 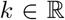 a free constant. Since the fixed point is stable, the last term vanishes in the long term. Let λ = *μ* + *iν*. The behavior of *α* greatly changes depending on whether *ν* is zero or not, i.e. whether the fixed point is a stable node or a stable focus. Let us start for the simple case, *ν* = 0. Then, at *t* → ∞,

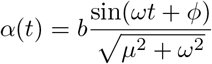

where *ϕ* = arctan(*μ/ω*). Therefore, these type of components always oscillate, but the amplitude of the oscillations decays as 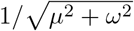. Hence, if the forcing frequency is too fast, or the stability too strong, then the induced oscillatory component becomes negligible.

Let’s turn now to the more interesting case of *ν* ≠ 0. Since we are considering a real system, there is always a pair of complex eigenvalues such that λ_±_ = *μ* ± *iν* associated to complex conjugate eigenvectors *α*_±_ (conjugate root theorem for polynomials with real coefficients). Therefore the dynamics of the real system is given by the real part of *α*_±_. We find that (for *t* → ∞),

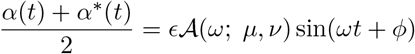

where

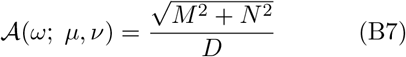

and *ϕ* = arctan(*N/M*), with

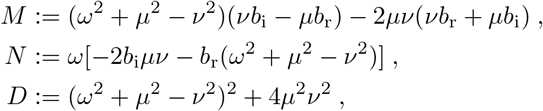

which corresponds to Eq. (24) in the main text.

## References

Aberra AS, Peterchev AV, Grill WM (2018) Biophysically realistic neuron models for simulation of cortical stimulation. Journal of Neural Engineering 15(6):066,023. https://doi.org/10.1088/1741-2552/aadbb1, URL http://stacks.iop.org/1741-2552/15/i=6/a=066023?key=crossref.c24583b2463818cf852344f5de358599

Agmon-Snir H, Segev I (1993) Signal delay and input synchronization in passive dendritic structures. Journal of Neurophysiology 70(5)

Augustin M, Ladenbauer J, Baumann F, et al (2017) Low-dimensional spike rate models derived from networks of adaptive integrate-and-fire neurons: Comparison and implementation. PLOS Computational Biology 13(6):e1005,545. https://doi.org/10.1371/journal.pcbi.1005545

Avermann M, Tomm C, Mateo C, et al (2012) Microcircuits of excitatory and inhibitory neurons in layer 2/3 of mouse barrel cortex. Journal of Neurophysiology 107(11):3116–34. https://doi.org/10.1152/jn.00917.2011

Bacci A, Rudolph U, Huguenard JR, et al (2003) Major differences in inhibitory synaptic transmission onto two neocortical interneuron subclasses. Journal of Neuroscience 23(29):9664–74. https://doi.org/10.1523/JNEUROSCI.23-29-09664.2003

Bartos M, Vida I, Frotscher M, et al (2002) Fast synaptic inhibition promotes synchronized gamma oscillations in hippocampal interneuron networks. Proceedings of the National Academy of Sciences 99(20):13,222–13,227. https://doi.org/10.1073/pnas.192233099

Bartos M, Vida I, Jonas P (2007) Synaptic mechanisms of synchronized gamma oscillations in inhibitory interneuron networks. Nature Reviews Neuroscience 8(1):45–56. https://doi.org/10.1038/nrn2044

Benayoun M, Cowan JD, van Drongelen W, et al (2010) Avalanches in a Stochastic Model of Spiking Neurons. PLoS Computational Biology 6(7):e1000,846. https://doi.org/10.1371/journal.pcbi.1000846

Bi H, di Volo M, Torcini A (2021) Asynchronous and coherent dynamics in balanced excitatory-inhibitory spiking networks. Frontiers in Systems Neuroscience 15. https://doi.org/10.3389/fnsys.2021.752261, URL https://www.frontiersin.org/article/10.3389/fnsys.2021.752261

Bikson M, Inoue M, Akiyama H, et al (2004) Effects of uniform extracellular DC electric fields on excitability in rat hippocampal slices in vitro. J Physiol 557(Pt 1):175–90

Brunel N, Hakim V (2008) Sparsely synchronized neuronal oscillations. Chaos: An Interdisciplinary Journal of Nonlinear Science 18(1):015,113. https://doi.org/10.1063/1.2779858

Buice MA, Cowan JD, Chow CC (2010) Systematic Fluctuation Expansion for Neural Network Activity Equations. Neural Computation 22(2):377–426. https://doi.org/10.1162/neco.2009.02-09-960

Buzsáki G, Wang XJ (2012) Mechanisms of Gamma Oscillations. Annual Review of Neuroscience 35(1):203–225. https://doi.org/10.1146/annurev-neuro-062111-150444

Byrne Á, O’Dea RD, Forrester M, et al (2020) Next-generation neural mass and field modeling. Journal of Neurophysiology 123(2):726–742. https://doi.org/10.1152/jn.00406.2019, pMID: 31774370

Byrne Á, Ross J, Nicks R, et al (2022) Mean-Field Models for EEG/MEG: From Oscillations to Waves. Brain Topography 35(1):36–53. https://doi.org/10.1007/s10548-021-00842-4

Cardin JA, Carlén M, Meletis K, et al (2009) Driving fast-spiking cells induces gamma rhythm and controls sensory responses. Nature 459(7247):663–667. https://doi.org/10.1038/nature08002

Carhart-Harris RL (2018) The entropic brain - revisited. Neuropharmacology https://doi.org/10.1016/j.neuropharm.2018.03.010.

Carlu M, Chehab O, Dalla Porta L, et al (2020) A mean-field approach to the dynamics of networks of complex neurons, from nonlinear Integrate-and-Fire to Hodgkin–Huxley models. Journal of Neurophysiology 123(3):1042–1051. https://doi.org/10.1152/jn.00399.2019

Chialvo DR (2004) Critical brain networks. Physica A: Statistical Mechanics and its Applications 340(4):756–765. https://doi.org/10.1016/j.physa.2004.05.064, URL https://doi.org/10.1016/j.physa.2004.05.064

Clusella P, Montbrió E (2022) in preparation, in preparation

Coombes S, Byrne Á (2019) Next generation neural mass models. In: Nonlinear dynamics in computational neuroscience. Springer, p 1–16

Deleuze C, Bhumbra GS, Pazienti A, et al (2019) Strong preference for autaptic self-connectivity of neocortical pv interneurons facilitates their tuning to *γ*-oscillations. PLOS Biology 17(9):e3000,419. https://doi.org/10.1371/journal.pbio.3000419

Destexhe A, Mainen ZF, Sejnowski TJ (1998) Kinetic models of synaptic transmission. In: Koch C, Segev I (eds) Methods in Neuronal Modeling, 2nd edn. MIT Press, Cambridge, MA., chap 1, p 1–25

Devalle F, Roxin A, Montbrió E (2017) Firing rate equations require a spike synchrony mechanism to correctly describe fast oscillations in inhibitory networks. PLOS Computational Biology 13(12)

Devalle F, Montbrió E, Pazó D (2018) Dynamics of a large system of spiking neurons with synaptic delay. Phys Rev E 98:042,214. https://doi.org/10.1103/PhysRevE.98.042214

Doedel EJ, Champneys AR, Dercole F, et al (2007) Auto-07p: Continuation and bifurcation software for ordinary differential equations

Dumont G, Gutkin B (2019) Macroscopic phase resetting-curves determine oscillatory coherence and signal transfer in inter-coupled neural circuits. PLoS Comput Biol 15(6)

Eeckman FH, J FW (1991) Asymmetric sigmoid non-linearity in the rat olfactory system. Brain Res 557(1–2):13–21

Ermentrout B (1994) Reduction of Conductance-Based Models with Slow Synapses to Neural Nets. Neural Computation 6(4):679–695. https://doi.org/10.1162/neco.1994.6.4.679

Ermentrout G, Bard DHTerman (2010) Mathematical Foundations of Neuroscience. Springer-Verlag New York

Eyal G, Verhoog MB, Testa-Silva G, et al (2018) Human cortical pyramidal neurons: From spines to spikes via models. Frontiers in Cellular Neuroscience 12. https://doi.org/10.3389/fncel.2018.00181, URL https://doi.org/10.3389/fncel.2018.00181

Forrester M, Crofts JJ, Sotiropoulos SN, et al (2020) The role of node dynamics in shaping emergent functional connectivity patterns in the brain. Network Neuroscience 4(2):467–483. https://doi.org/10.1162/netn_a_00130, URL https://doi.org/10.1162/netn_a_00130

Fourcaud-Trocmé N, Hansel D, van Vreeswijk C, et al (2003) How spike generation mechanisms determine the neuronal response to fluctuating inputs. Journal of Neuroscience 23(37):11,628–11,640. https://doi.org/10.1523/JNEUROSCI.23-37-11628.2003, URL https://www.jneurosci.org/content/23/37/11628

Freeman WJ (1972) Linear Analysis of the Dynamics of Neural Masses. Annual Review of Biophysics and Bioengineering 1(1):225–256. https://doi.org/10.1146/annurev.bb.01.060172.001301

Freeman WJ (1975) Mass Action in the Nervous System. New York: Academic Press

Freeman WJ (1987) Simulation of chaotic EEG patterns with a dynamic model of the olfactory system. Biol Cybern 56(2-3):139–50

Galan A (2021) Realistic modeling of neocortical neurons and electric field effects under direct current stimulation. Master’s thesis, Elite Master Program in Neuroengineering, Department of Electrical and Computer Engineering, Technical University of Munich

Gerstner W (1995) Time structure of the activity in neural network models. Physical Review E 51(1):738–758. https://doi.org/10.1103/PhysRevE.51.738

Goldobin DS, di Volo M, Torcini A (2021) Reduction methodology for fluctuation driven population dynamics. Phys Rev Lett 127:038,301. https://doi.org/10.1103/PhysRevLett.127.038301, URL https://link.aps.org/doi/10.1103/PhysRevLett.127.038301

Grimbert F, Faugeras O (2006) Analysis of Jansen’s model of a single cortical column. INRIA RR-5597:34

Jang HJ, Cho KH, Park SW, et al (2010) The development of phasic and tonic inhibition in the rat visual cortex. Korean J Physiol Pharmacol 14:299–405

Jansen BH, Rit VG (1995) Electroencephalogram and visual evoked potential generation in a mathematical model of coupled cortical columns. Biol Cybern 73(4):357–66

Jansen BH, Zouridakis G, Brandt ME (1993) A neurophysiologically-based mathematical model of flash visual evoked potentials. Biol Cybern 68(3):275–83

Jedynak M, Pons AJ, Garcia-Ojalvo J, et al (2017) Temporally correlated fluctuations drive epileptiform dynamics. NeuroImage 146:188–196

Karnani MM, Jackson J, Ayzenshtat I, et al (2016) Cooperative subnetworks of molecularly similar interneurons in mouse neocortex. Neuron 90(1):86–100. https://doi.org/10.1016/j.neuron.2016.02.037

Kay LM (2018) The physiological foresight in Freeman’s work. J Conscious Stud 25(1–2):50–63

Koch C, Segev I (eds) (2003) Methods in neuronal modeling, 2nd edn. Computational Neuroscience Series, Bradford Books, Cambridge, MA

Kunze T, Hunold A, Haueisen J, et al (2016) Transcranial direct current stimulation changes resting state functional connectivity: A large-scale brain network modeling study. NeuroImage 140:174–187. https://doi.org/10.1016/j.neuroimage.2016.02.015, URL http://dx.doi.org/10.1016/j.neuroimage.2016.02.015

Laing CR (2015) Exact Neural Fields Incorporating Gap Junctions. SIAM Journal on Applied Dynamical Systems 14(4):1899–1929. https://doi.org/10.1137/15M1011287

Latham PE, Richmond BJ, Nelson PG, et al (2000) Intrinsic Dynamics in Neuronal Networks. I. Theory. Journal of Neurophysiology 83(2):808–827. https://doi.org/10.1152/jn.2000.83.2.808

Lopez-Sola E, Sanchez-Todo R, Lleal È, et al (2021) A personalizable autonomous neural mass model of epileptic seizures. BiorXiv https://doi.org/10.1101/2021.12.24.474090, URL https://doi.org/10.1101/2021.12.24.474090

Merlet I, Birot G, Salvador R, et al (2013) From Oscillatory Transcranial Current Stimulation to Scalp EEG Changes: A Biophysical and Physiological Modeling Study. PLoS ONE 8(2):1–12. https://doi.org/10.1371/journal.pone.0057330

Molaee-Ardekani B, Benquet P, Bartolomei F, et al (2010) Computational modeling of high-frequency oscillations at the onset of neocortical partial seizures: from ‘altered structure’ to ‘dysfunction’. Neuroimage 52(3):1109–22

Montbrió E, Pazó D (2020) Exact mean-field theory explains the dual role of electrical synapses in collective synchronization. Phys Rev Lett 125:248,101. https://doi.org/10.1103/PhysRevLett.125.248101, URL https://link.aps.org/doi/10.1103/PhysRevLett.125.248101

Montbrió E, Pazó D, Roxin A (2015) Macroscopic description for networks of spiking neurons. Phys Rev X 021028

Neske GT, Patrick SL, Connors BW (2015) Contributions of diverse excitatory and inhibitory neurons to recurrent network activity in cerebral cortex. Journal of Neuroscience 35(3):1089–1105. https://doi.org/10.1523/JNEUROSCI.2279-14.2015

Oláh S, Komlósi G, Szabadics J, et al (2007) Output of neurogliaform cells to various neuron types in the human and rat cerebral cortex. Frontiers in Neural Circuits 1. https://doi.org/10.3389/neuro.04.004.2007

Ostojic S, Brunel N (2011) From Spiking Neuron Models to Linear-Nonlinear Models. PLoS Computational Biology 7(1):e1001,056. https://doi.org/10.1371/journal.pcbi.1001056

Pazó D, Montbrió E (2016) From quasiperiodic partial synchronization to collective chaos in populations of inhibitory neurons with delay. Phys Rev Lett 116:238,101. https://doi.org/10.1103/PhysRevLett.116.238101, URL https://link.aps.org/doi/10.1103/PhysRevLett.116.238101

Pereira U, Brunel N (2018) Attractor Dynamics in Networks with Learning Rules Inferred from In Vivo Data. Neuron 99(1):227–238.e4. https://doi.org/10.1016/j.neuron.2018.05.038

Pietras B, Devalle F, Roxin A, et al (2019) Exact firing rate model reveals the differential effects of chemical versus electrical synapses in spiking networks. Phys Rev E 100:042,412

Pods J, Schönke J, Bastian P (2013) Electrodiffusion models of neurons and extracellular space using the poisson-nernst-planck equations— numerical simulation of the intra- and extracellular potential for an axon model. Biophysical Journal 105:242–254

Pons AJ, Cantero JL, Atienza M, et al (2010) Relating structural and functional anomalous connectivity in the aging brain via neural mass modeling. Neuroimage 52(3):848–861

Povysheva NV, Zaitsev AV, Kröner S, et al (2007) Electrophysiological differences between neurogliaform cells from monkey and rat prefrontal cortex. Journal of Neurophysiology 97(2). https://doi.org/10.1152/jn.00794.2006

Ratas I, Pyragas K (2016) Macroscopic self-oscillations and aging transition in a network of synaptically coupled quadratic integrate-and-fire neurons. Physical Review E 94(3):032,215. https://doi.org/10.1103/PhysRevE.94.032215

Ratas I, Pyragas K (2018) Macroscopic oscillations of a quadratic integrate-and-fire neuron network with global distributed-delay coupling. Phys Rev E 98:052,224. https://doi.org/10.1103/PhysRevE.98.052224, URL https://link.aps.org/doi/10.1103/PhysRevE.98.052224

Ratas I, Pyragas K (2019) Noise-induced macroscopic oscillations in a network of synaptically coupled quadratic integrate-and-fire neurons. Physical Review E 100(5):052,211. https://doi.org/10.1103/PhysRevE.100.052211

Rauch A, Camera GL, Lüscher HR, et al (2003) Neocortical pyramidal cells respond as integrate-and-fire neurons to in vivo–like input currents. J Neurphysiol 90:1598–1612

Ruffini G, Lopez-Sola E (2022) AIT foundations of structured experience

Ruffini G, Wendling F, Merlet I, et al (2013) Transcranial Current Brain Stimulation (tCS): Models and Technologies. IEEE Transactions on Neural Systems and Rehabilitation Engineering 21(3):333–345

Ruffini G, Wendling F, Sanchez-Todo R, et al (2018) Targeting brain networks with multichannel transcranial current stimulation (tcs). Current Opinion in Biomedical Engineering

Ruffini G, Salvador R, Tadayon E, et al (2020) Realistic modeling of mesoscopic ephaptic coupling in the human brain. PLoS Comput Biol

Sanchez-Todo R, Salvador R, Santarnecchi E, et al (2018) Personalization of hybrid brain models from neuroimaging and electrophysiology data. BioRxiv 00(0):1–35. https://doi.org/10.1101/461350, URL https://www.biorxiv.org/content/10.1101/461350v1

Seay M, Natan RG, Geffen MN, et al (2020) Differential short-term plasticity of PV and SST neurons accounts for adaptation and facilitation of cortical neurons to auditory tones. Journal of Neuroscience 40(48):9224–9235. https://doi.org/10.1523/JNEUROSCI.0686-20.2020

Lopes da Silva F, Hoek A, Smits H, et al (1974) Model of brain rhythmic activity: the alpha rhythm of the thalamus. Kybernetik 15(1):27–37

Lopes da Silva F, A vR, Barts P, et al (1976) Model of neuronal populations: the basic mechanism of rhythmicity. Prog Brain Res 45

Stefanovski L, Triebkorn P, Spiegler A, et al (2019) Linking molecular pathways and large-scale computational modeling to assess candidate disease mechanisms and pharmacodynamics in alzheimer’s disease. Front Comput Neurosci

Taher H, Torcini A, Olmi S (2020) Exact neural mass model for synaptic-based working memory. PLOS Computational Biology 16(12):1–42. https://doi.org/10.1371/journal.pcbi.1008533, URL https://doi.org/10.1371/journal.pcbi.1008533

Taher H, Avitabile D, Desroches M (2022) Bursting in a next generation neural mass model with synaptic dynamics: A slow–fast approach. Nonlinear Dynamics https://doi.org/10.1007/s11071-022-07406-6

Tiesinga P, Sejnowski TJ (2009) Cortical Enlightenment: Are Attentional Gamma Oscillations Driven by ING or PING? Neuron 63(6):727–732. https://doi.org/10.1016/j.neuron.2009.09.009

Traub RD, Spruston N, Soltesz I, et al (1998) Gamma-frequency oscillations: A neuronal population phenomenon, regulated by synaptic and intrinsic cellular processes, and inducing synaptic plasticity. Progress in Neurobiology 55(6):563–575. https://doi.org/10.1016/S0301-0082(98)00020-3

Van Vreeswijk C, Abbott LF, Bard Ermentrout G (1994) When inhibition not excitation synchronizes neural firing. Journal of Computational Neuroscience 1(4):313–321. https://doi.org/10.1007/BF00961879, URL https://doi.org/10.1007/BF00961879

Vázquez-Rodríguez B, Avena-Koenigsberger A, Sporns O, et al (2017) Stochastic resonance at criticality in a network model of the human cortex. Scientific Reports 7(1). https://doi.org/10.1038/s41598-017-13400-5, URL https://doi.org/10.1038/s41598-017-13400-5

di Volo M, Torcini A (2018) Transition from asynchronous to oscillatory dynamics in balanced spiking networks with instantaneous synapses. Phys Rev Lett 121:128,301. https://doi.org/10.1103/PhysRevLett.121.128301, URL https://link.aps.org/doi/10.1103/PhysRevLett.121.128301

Wendling F, Chauvel P (2008) Transition to Ictal Activity in Temporal Lobe Epilepsy: Insights From Macroscopic Models. Computational Neuroscience in Epilepsy pp 356–386. https://doi.org/10.1016/B978-012373649-9.50026-0

Wendling F, Bartolomei F, Bellanger JJ, et al (2002) Epileptic fast activity can be explained by a model of impaired GABAergic dendritic inhibition. Eur J Neurosci 15(9):1499–508

Whittington M, Traub R, Kopell N, et al (2000) Inhibition-based rhythms: Experimental and mathematical observations on network dynamics. International Journal of Psychophysiology 38(3):315–336. https://doi.org/10.1016/S0167-8760(00)00173-2

Whittington MA, Traub RD, Jefferys JGR (1995) Synchronized oscillations in interneuron networks driven by metabotropic glutamate receptor activation. Nature 373(6515):612–615. https://doi.org/10.1038/373612a0

Wilson HR, Cowan JD (1972) Excitatory and Inhibitory interactions in localized populations of model neurons. Biophysical Journal 12(1):1–24. https://doi.org/10.1016/S0006-3495(72)86068-5, URL http://dx.doi.org/10.1016/S0006-3495(72)86068-5

Zaitsev AV, Povysheva NV, Gonzalez-Burgos G, et al (2012) Electrophysiological classes of layer 2/3 pyramidal cells in monkey prefrontal cortex. Journal of Neurophysiology 108(2):595–609. https://doi.org/10.1152/jn.00859.2011

Zimmern V (2020) Why brain criticality is clinically relevant: A scoping review. Front Neural Circuits 14

